# Whole blood and hypertonic saline resuscitation improve the outcomes of controlled hemorrhagic shock combined with penetrating brain injury in rat

**DOI:** 10.1101/448266

**Authors:** Marina Weissmann, Dalit E. Dar, Michael M. Krausz

## Abstract

One of the most frequent injury patterns leading to early death of polytraumatized patients is hemorrhagic shock combined with traumatic brain injury. Currently, there is no agreement on the best treatment protocol of such injury. This study was aimed to compare the outcomes of resuscitation with various volumes of hypertonic saline (HTS), whole blood (WB) and fresh frozen plasma (FFP) in a rat model of penetrating brain injury combined with controlled hemorrhagic shock in order to look for optimal treatment. Anesthetized rats were intubated and ventilated. 30% of circulating blood volume was withdrawn following open brain injury, and intravenous resuscitation with various volumes of HTS, WB and FFP (3, 9, and 18 ml/kg) was administered. Blood samples were collected during the experiment for measurements of lactate and base excess levels, and before sacrifice for neutrophil counts. Twenty four and 48 hours after the injury, rat’s neurological status was examined. Rats were, then, sacrificed and pathological studies of brain and lung sections were performed. Regardless of fluid type, resuscitation improved the mean arterial pressure and lactate levels in accordance to fluid’s volume. However, base excess was improved only when rats were treated with WB and FFP. Improvement in behavioral performance of the rats was observed when treated with 3 and 9 ml/kg HTS or with 18 ml/kg WB or FFP. Blood and lung neutrophils were reduced in rats treated with 9 or 18 ml/kg WB. Mortality rates were reduced in rats administered with 9 and 18 ml/kg of HTS or WB. Taken together, resuscitation with WB at 9-18ml/kg or HTS at 9ml/kg is optimal for treatment of combined injury.

## Introduction

Multiple trauma is among the leading causes of death and permanent disability during injuries [1]. One of the most frequent injury patterns leading to early death of polytraumatized patients is hemorrhagic shock combined with traumatic brain injury (BI) [2]. Hemorrhagic shock is characterized by decrease in mean arterial pressure (MAP), hypoxia, metabolic acidosis and coagulopathy [3]. Traumatic BI is increased by injuries such as: hypoxia and hypovolemia, which exacerbate brain ischemia and raise the probability of mortality [4].

According to Advanced Trauma Life Support (ATLS) recommendations, fluid resuscitation should begin with 2L of crystalloid solution. If the patient does not respond to initial crystalloid administration, blood products should be added. The recommended ratio between packed red blood cells (PRBCs) and fresh frozen plasma (FFP) is 1:1, starting with administration of PRBCs to ensure adequate oxygen delivery, and FFP and platelets are added at a later stage [5-9]. This resuscitation protocol raises two issues that should be addressed: First, aggressive crystalloid administration to patients with BI may lead to brain swelling due to transition of fluids into the interstitial space, which in turn may aggravate the secondary BI [10, 11]. Second, according to Harhanghi et al., one out of three patients with traumatic BI arrives at the hospital with signs of coagulopathy due to dilution of the blood by resuscitation fluids and consumption of coagulation factors [12]. This coagulopathy correlates with increased incidence of secondary brain injuries [13] and may contribute to increased mortality [14, 15]. Thus, several previous studies suggested that the use of earlier administration of FFP and higher ratio between PRBCs and FFP may be beneficial in acute coagulopathic patients [9, 16, 17].

Animal studies using controlled hemorrhagic shock (CHS) combined with BI have shown that HTS treatment was beneficial in comparison to isotonic crystalloids in reducing intracranial pressure and brain edema and in behavioral and neuronal recovery [18-22]. BI models used in these studies were models of blunt brain injury. To our knowledge, no study tested the response of HTS or FFP on CHS combined with penetrating BI (e.g. gunshot wounds, missiles). This mechanism of impact, although less prevalent than blunt injury, has much worse prognosis due to distortion of tissues that lie directly in the path of the projectile, increased chance of hemorrhage from the open wound and increased probability of infections in the brain tissue [23].

The goal of the present study was to search for the optimal treatment by comparing HTS, FFP, and whole blood (WB) on a rat model of penetrating BI combined with CHS, using various resuscitation volumes.

## Materials and Methods

### Animals

Male Lewis rats weighing 300-350 gram (Harlan Laboratories, Rehovot, Israel) were maintained in the animal facilities of the Faculty of Medicine of the Technion on a 12:12 light-dark cycle with free access to standard laboratory food and water. All animal experiments accorded with the requirements of the Technion’s Care and Use Committee, which completely accord with the National Institutes of Health Guide for Care and Use of Laboratory Animals. All surgery was performed under sodium pentobarbital anesthesia, and all efforts were made to minimize suffering.

Rats were divided into 13 groups: sham operated animals (n=8), animals subjected to BI (n=12) or CHS (n=8) and animals subjected to combined injury (n=16). In the other nine groups, animals were subjected to combined injury and treated for one hour with WB, FFP or HTS in three different volumes (n=10-14).

### Experimental setting

Rats were anesthetized with a mixture of Pental (50 mg/Kg) and Fentanyl (0.65 μg/Kg) administered intraperitoneally, endotracheal intubated using dim cut venflon tube and ventilated with fresh air (2.5ml volume, 110-120 BPM), [Harvard Apparatus, Holliston, USA]. The right and left femoral arteries and left femoral vein were cannulated using polyethylene tubing. The right arterial line was connected to a calibrated pressure transducer and to a controlled data acquisition system [Trasducer MLT0699, Amplifier ML224, PowerLab 4/30 ML866, ADInstruments, Sydney, Australia] for continuous blood pressure monitoring. The left femoral artery was used for blood withdrawal and the vein for intravenous fluid infusion. At the end of the surgical procedure, the tubing was removed, and the femoral arteries and vein were ligated. BI was induced using the “dynamic cortical deformation” model, as previously described [24, 25]. Briefly, after scalp insicion, 5 mm diameter burr hole was drilled in the left parietal region, the dura was removed and negative pressure of 300 torr (40kPa) was applied to the cortical surface for 10 sec. The skin was thereafter sutured. Five minutes after induction of BI, 30% of the circulating blood volume was withdrawn through the left femoral artery. MAP was reduced from 150 to 40mmHg and was maintained at this level for 40 minutes. 5 minutes after the end of hemorrhage, resuscitation was administered using syringe pump [Graseby Medical Ltd., Watford, UK] for 1 hour. Blood samples were collected into heparinized glass capillary tubes [Instrumentation Laboratory, Lexington, USA] at 0 (before BI), 30 (after BI), and 85 (after CHS) minutes, as well as 100, 115 (15 & 30 minutes after the beginning of resuscitation accordingly) and 145 minutes (end of treatment). Blood lactate, base excess, hematocrit and electrolyte levels were determined by GEM Premier 3000 [Instrumentation Laboratory, Lexington, USA]. MAP was computed from the arterial tracing. Fig 1A shows a schematic representation of the experiment timeline.

**Fig.1.**
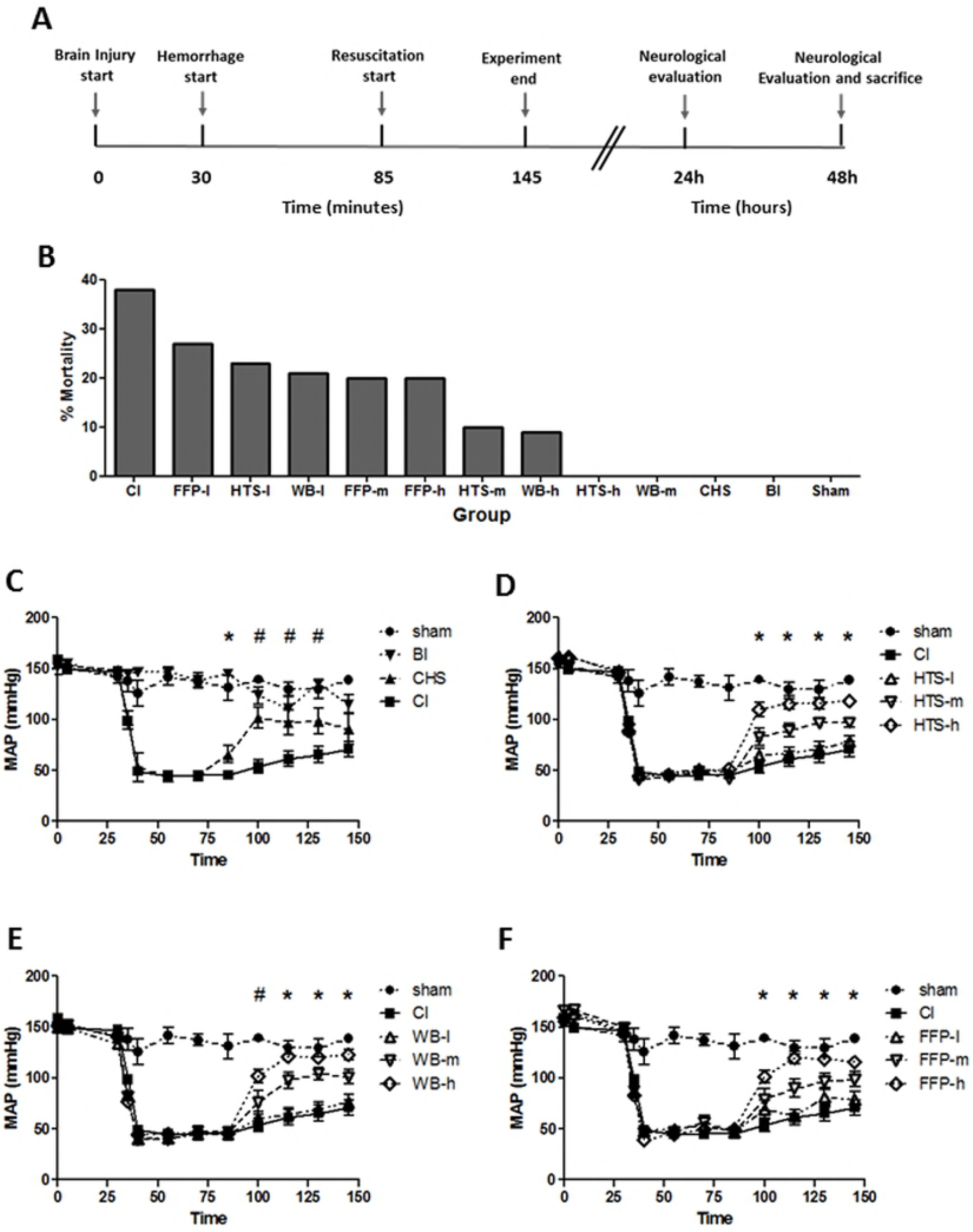
Mean arterial pressure. Rats were anesthetized and intubated, and cannulas were inserted into femoral arteries and vein. Experimental design is represented in **(A)** and mortality rate during the first 24 hours after the injury in **(B)**. Mean arterial pressure was computed from the arterial tracing and is shown in control groups **(C)** * P<0.05 CI, CHS vs. sham, BI; # P< 0.05 CI vs. CHS, and in resuscitation groups **(D)-(F)**. **(D)** Hypertonic Saline (HTS) resuscitation: * P<0.05 CI vs. HTS-m, HTS-h. **(E)** Whole Blood (WB) resuscitation: # P<0.05 CI vs. WB-h; * P<0.05 CI vs. WB-m, WB-h. **(F)** Fresh Frozen Plasma (FFP) resuscitation: * P<0.05 CI vs. FFP-h. **BI**-Brain Injury, **CHS**-Controlled Hemorrhagic Shock, **CI**-combined injury, **l**-low (3 ml/kg), **m**-moderate (9 ml/kg), **h**-high (18 ml/kg) volume.

### Resuscitation fluids

A group of animals served as blood donors. Lewis rats were used therefore blood typing was not necessary before transfusion. Blood was withdrawn through the carotid artery into a sterile tube containing citrate phosphate dextrose-adenine. Whole blood was stored at 4°C. For FFP preparation, blood was centrifuged at 3,300 RPM, the supernatant was separated and stored at-20°C. HTS (7.5%) was freshly prepared for each group. Three volumes were used with each type of resuscitation fluid: 3 ml/kg (low), 9 ml/kg (moderate) or 18 ml/kg (high).

### Neurological status evaluation

Twenty four and 48 hours after surgery, neurological state of the rat was assessed by the neurological severity score (NSS) grading system, as previously described [25]. Points were assigned for alteration of motor function and behavior so that a maximal score of 25 represents severe neurological dysfunction whilst a score of 0 indicates an intact neurological condition (opposite to the human Glasgow Coma Scale).

### Flow cytometry

After completion of functional testing, rats were sacrificed, and blood samples were immediately taken from vena cava, before the beginning of trans-cardial perfusion. Blood samples were lysed with BD FACS Lysing Solution [BD Biosciences, San Jose, USA], washed with PBS supplemented with 3mM EDTA and 1% FCS, and resuspended with 0.5% PFA. Samples were analyzed by the Cyan ADP 3 laser Flow cytometer [Beckman Coulter Inc, Nyon, Switzerland] on the next day. Leucocytes were gated according to the typical forward and side scatter profiles.

### Pathological examination

Transcardial perfusion with 120ml of heparinized saline followed by 120 ml of 4% formaldehyde was conducted, as previously described [25]. The brain and a section of the right lung lobe were removed, fixated with 4% formaldehyde, and embedded in paraffin, for histological examination.

Brain tissues were stained with NeuN antibody, as previously described [25]. Neutrophil staining of the lungs was performed using the Naphtol AS-D chloroacetate/hematoxilin kit [Sigma, St. Louis, USA] according to the manufacturer’s instructions. The amount of stained neurons and neutrophils for each animal was counted and averaged from 5 fields on each slide.

### Statistical tests

All results are expressed as the mean ±SEM. Analyses were performed using Prism GraphPad software. Groups were compared using one-way ANOVA and *t* tests analyses. P<0.05 was considered significant.

## Results

### Mortality

The first 24 hours after the injury were critical for the survival of the rats. During this period, while no mortality was observed in groups subjected to penetrating BI or CHS alone, the combination of both resulted in mortality rate of 38% (CI, Fig 1B). Treatment with low volume of resuscitation did not improve mortality in neither group, however treatment with moderate and high volumes of WB and HTS significantly reduced mortality rate to 10% and lower (Fig 1B).

### Hemodynamic and metabolic measurements

10 minutes after the beginning of controlled bleeding, blood pressure of rats dropped to approximately 40mmHg and was manually maintained at this level for additional 40 minutes. Significant improvement in blood pressure of CHS rats, compared to CI rats, was observed 15, 30 and 45 (P< 0.05) minutes after hemorrhage stage (Fig 1C). Treatment with low volume of resuscitation fluids did not result in significant improvement in MAP at the end of 1 hour resuscitation. However, MAP was elevated to approximately 100mmHg in moderate and 120mmHg in high (P<0.05) resuscitation groups, regardless of the resuscitation fluid used (Figs 1D-F). Importantly, blood pressure of 75% of rats that died during the study was lower than 80mmHg at the end of 1 hour observation/ resuscitation period.

Hemorrhage resulted in elevation of lactate level to 4.0±0.4 and 4.7±0.4 mmol/L in CHS and CI groups respectively (P<0.05 CI, CHS vs. sham, BI), and reduction in BE level to −1.7±0.5 mmol/L and −4.5±0.8 mmol/L respectively (P<0.05 CI, CHS vs. sham, BI). During the observation period, lactate and BE levels of the CHS group gradually returned to the normal values, while lactate and BE levels of CI rats showed only a slight improvement (P<0.05 CI vs. CHS, Figs 2A and 3A). In resuscitation groups, improvement in lactate level was in correlation with resuscitation volume and appeared parallel to the improvement in MAP. In rats resuscitated with moderate volume of WB, or high volumes of WB, FFP or HTS, lactate levels returned to normal by the end of resuscitation period (P<0.05 CI vs. HTS-h, FFP-h, WB-m, WB-h, Figs 2B-D). Rats treated with WB and FFP had improved BE levels by the end of resuscitation period (P<0.05, Figs 3C and D), however, only low volume HTS treatment resulted in improved BE levels (P<0.05, Fig 3B).

**Fig.2.**
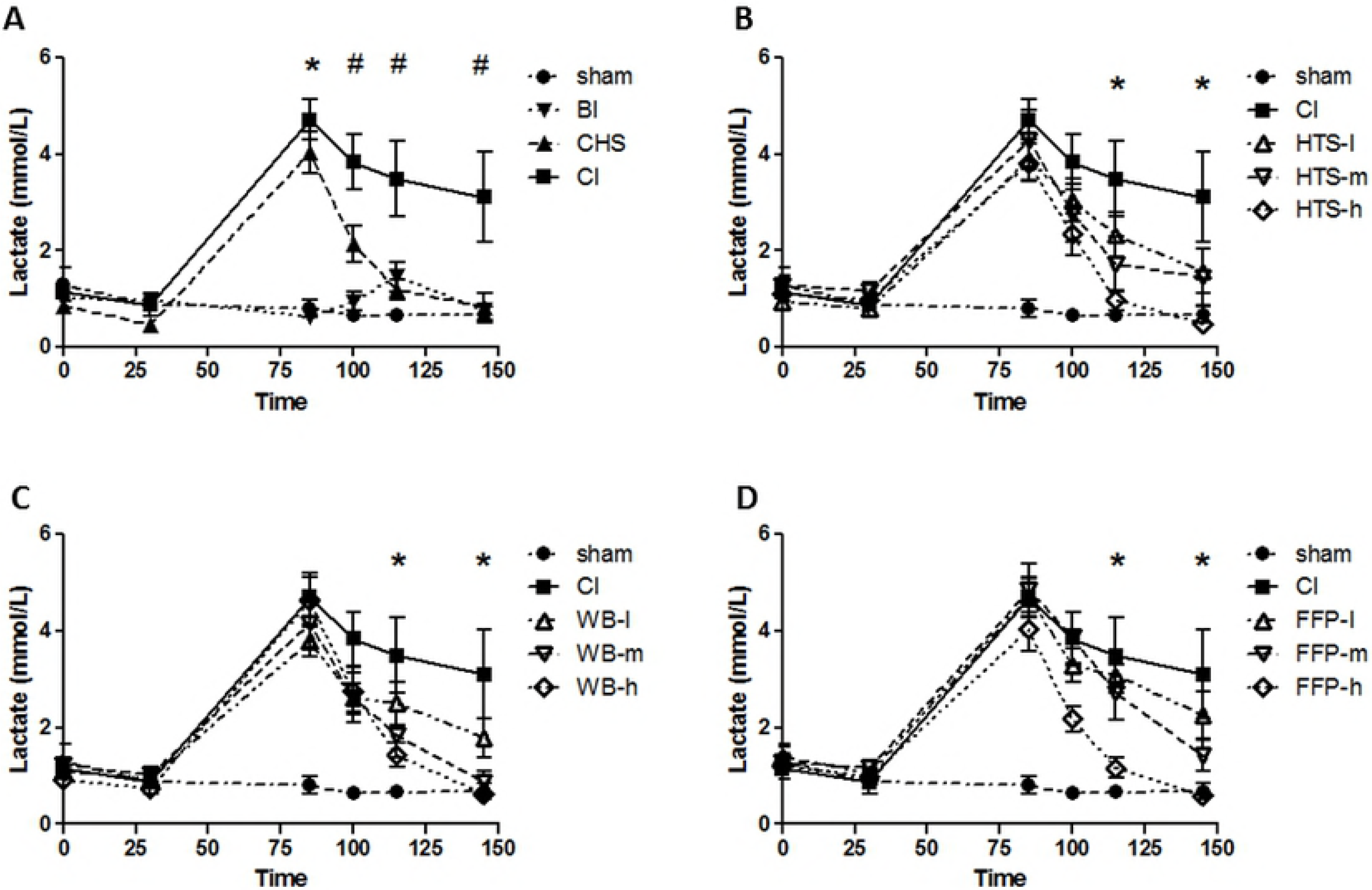
Blood lactate. Arterial blood samples were collected at different time-points during the experiment and lactate level in rat’s blood was measured. **(A)** Control groups: * P<0.05 CI, CHS vs. sham, BI; * P< 0.05 CI vs. CHS. **(B)** Hypertonic Saline (HTS) resuscitation, **(C)** Whole Blood (WB) resuscitation, and **(D)** Fresh Frozen Plasma (FFP) resuscitation. * P<0.05 CI vs. HTS-m, HTS-h, WB-m, WB-h, FFP-h. **BI**-Brain Injury, **CHS**-Controlled Hemorrhagic Shock, **CI**-combined injury, l-low (3 ml/kg), **m**-moderate (9 ml/kg), **h**-high (18 ml/kg) volume.

**Fig.3.**
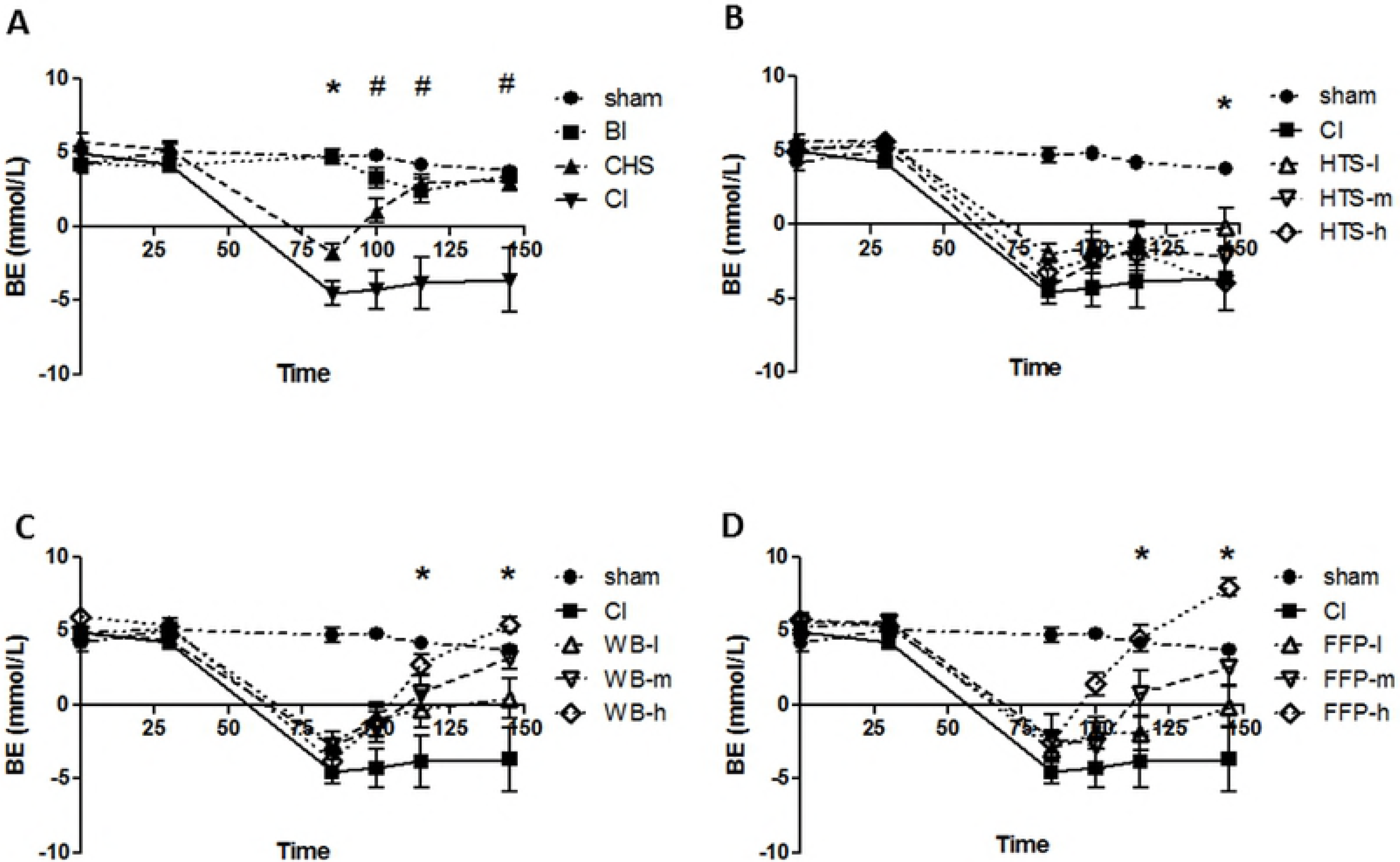
Base Excess in blood. Blood acidity at different time-points is represented by base excess, which was calculated based on the pH and pC0_2_ measurements. **(A)** Control groups: * P<0.05 CI, CHS vs. sham, BI; # P< 0.05 CI vs. CHS. **(B)** Hypertonic Saline (HTS), **(C)** Whole Blood (WB), and **(D)** Fresh Frozen Plasma (FFP) resuscitation. * P<0.05 CI vs. HTS-l, WB-m, WB-h, FFP-m, FFP-h. **BI**-Brain Injury, **CHS**-Controlled Hemorrhagic Shock, **CI**-combined injury, l-low (3 ml/kg), **m**-moderate (9 ml/kg), **h**-high (18 ml/kg) volume.

### Brain injury evaluation

Twenty four and 48 hours after the injury, rats’ neurological status was tested using the neurological severity score (NSS) grading system. Rats that were subjected to combined injury had higher severity score compared to rats subjected to BI alone, measured 24 (score of 6.6±1.3 compared with 3.3±1, P <0.05) and 48 (score of 5.4±1.2 compared with 1.5±0.8, P< 0.005) hours after brain trauma (Fig 4A). NSS of sham and CHS rats was “0”, presenting no neurological dysfunction. Improved NSS scores were observed in rats treated with high volume of blood fluids, and limited and moderate volumes of HTS, 48 hours after the injury (P<0.05 CI vs. WB-h, FFP-h, HTS-l, HTS-m Figs 4B-D).

**Fig.4.**
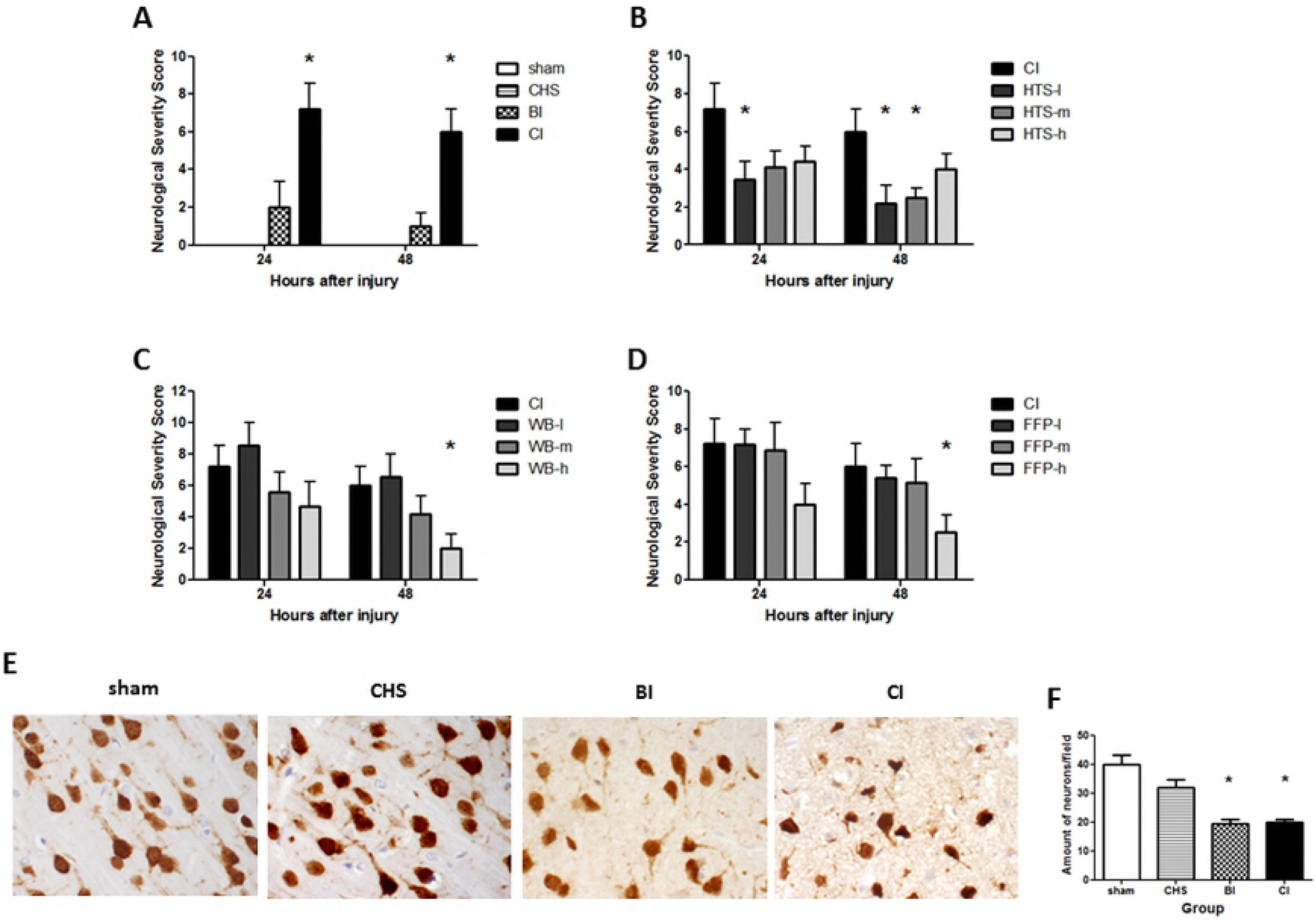
Evaluation of brain injury extent. 24 and 48 hours after the injury, neurological damage was assessed using a Neurological Severity Score test, which measures rat’s ability to move, balance on beam walking, and rat’s reflects. Scores are measured from 0 – no neurological dysfunction to 25 – complete inability to move. Test scores are shown in figures **(A)-(D)**. **(A)** Control groups, **(B)** Hypertonic Saline (HTS) resuscitation, **(C)** Whole Blood (WB) resuscitation, **(D)** Fresh Frozen Plasma (FFP) resuscitation. * P<0.05 CI vs. HTS-l, HTS-m, WB-h, FFP-h. 48 hours after the experiment, rats were sacrificed and brain pathology was examined applying anti-neuronal staining in the lesion area. Representative histological sections from cortical perilesional area are shown in **(E)**, and neuron counts are shown in **(F)**. * P<0.05 CI, BI vs. sham, CHS. **BI**-Brain Injury, **CHS**-Controlled Hemorrhagic Shock, **CI**-combined injury, **l**-low (3 ml/kg), **m**-moderate (9 ml/kg), h-high (18 ml/kg) volume.

Pathological examination of brain tissues which lay adjacent to the damaged area revealed that the shape and size of neurons in these areas were changed: neuronal cells were shrunk, with small amorphous nuclei in comparison to the round shape in healthy brain tissue (Fig 4E). The amount of neurons in this area was significantly lower in BI and CI groups, compared to the parallel area in sham and CHS brains (P<0.05 BI, CI vs. CHS, sham, Fig 4F). The amount and morphology of neurons was not improved by either resuscitation protocol used.

### Immunological response

Forty eight hours after the injury, the amount of neutrophils in the circulation was measured. Significantly high neutrophil counts in the blood were found in BI, CHS, and CI groups, compared to sham operated rats (P<0.05, Fig 5A). Since elevated neutrophil counts in the circulation can lead to invasion of these neutrophils into the lungs, we examined the lung tissue by immunohistochemical staining. As we suspected, high amount of neutrophils invaded the pulmonary tissue after the trauma (P<0.05 BI, CHS, CI vs. sham, Figs 5C and D). Additionally, lung tissue of rats subjected to combined injury was thicker with swollen alveolar walls, compared to lung tissue of sham rats. Treatment with moderate or high volumes of WB resulted in lower neutrophil counts in the circulation (P<0.05, Fig 5B), and decreased neutrophil recruitment to the lungs (P<0.05, Figs 5C and D).

**Fig.5.**
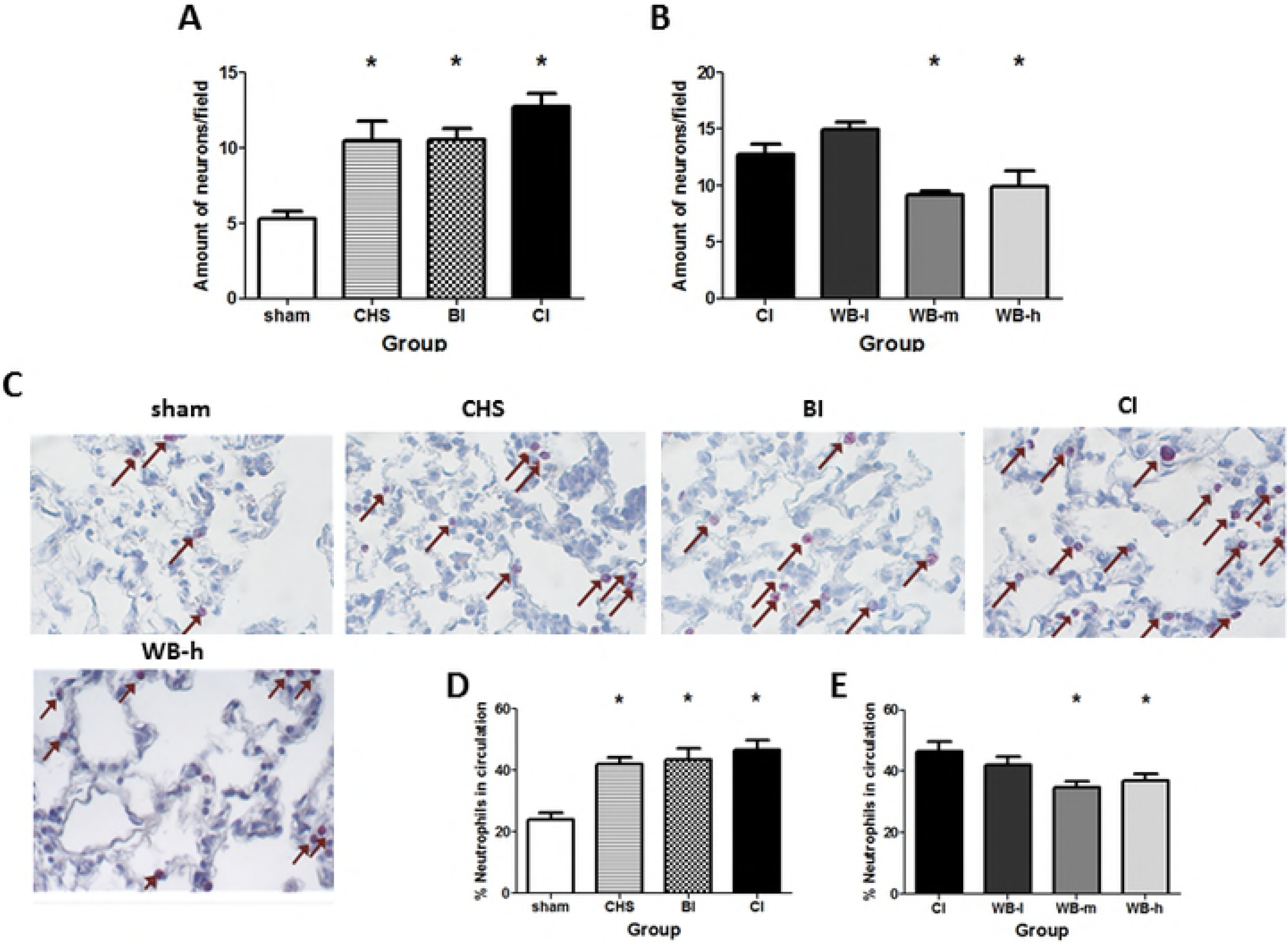
Inflammatory response after combined injury in the rat. Before sacrifice, blood samples were collected from vena cava and flow cytometry analysis of the samples was performed. Leukocytes were gated according to standard forward and side scatter characteristics, and amount of neutrophils was counted. Shown here are neutrophil counts in control groups **(A)** * P<0.05 sham vs. BI, CHS, CI, and in Whole Blood (WB) treated groups **(B)** *P<0.05 CI vs. WB-m, WB-h. Lung sections were stained for neutrophils, to evaluate the extent of neutrophil infiltration into the lung. **(C)** Representative histological sections of lungs. The arrows point on neutrophil cells. The amount of neutrophils per field was counted in each section is shown in **(D)** control groups *p<0.05 sham vs. BI, CHS, CI, and in **(E)** WB treated groups, *P<0.05 CI vs. WB-m, WB-h. **BI**-Brain Injury, **CHS**-Controlled Hemorrhagic Shock, **CI**-combined injury, l-low (3 ml/kg), **m**-moderate (9 ml/kg), **h**-high (18 ml/kg) volume.

## Discussion

The goal of this study was to establish resuscitation protocol for treatment of penetrating BI combined with controlled hemorrhage. Using three types of resuscitation fluids-HTS, WB, and FFP, we aimed to determine the best resuscitation fluid and best resuscitation volume to be used for beneficial neurological outcome.

Since most of the studies on BI use the fluid percussion injury model, which imitates blunt injury, we chose to use a model of focal penetrating BI, the prognosis of which is much worse clinically [23]. The method of dynamic cortical deformation was originally described by Shreiber [24], and was characterized in respect to the extent of the damage. It was shown that the impact area was clearly delineated and differentiated from spared surrounding brain tissue [24, 26], and that the amount of apoptotic cells and inflammatory markers in the perilesional area were increased. In our previous study, using this method, we did not see changes in the neurological damage between rats subjected to isolated BI or those subjected to combined injury, and no mortality was observed in the CI group [25]. Therefore, in the present study, we improved the model by reducing the extent of BI, adding mechanical ventilation, and prolonging the bleeding period. As a result, we were able to show that the combination of HS with BI exacerbated hemodynamic and metabolic measurements caused by HS, and intensified the cognitive and coordinative disabilities of the rat caused by BI, as well as elevating mortality rate to 38%.

Hemorrhagic shock resulted in metabolic acidosis, characterized by low BE and high lactate levels in the blood. Improvement in MAP and lactate levels was observed in volume-dependent manner in all resuscitation groups, however, BE levels were not improved in rats treated with moderate or high HTS volumes, despite the decrease in lactic acid levels in the blood. This effect of HTS administration was explained previously by sharp elevation in hydrochloric acid in the plasma, which corresponded with the steep rise in blood sodium levels observed in rats treated with moderate or high volumes of HTS **(S1 Fig)**, thus resulting in development of hyperchloremic acidosis [27].

The normalization of physiological signs is crucial to survival after trauma, thus we assumed that faster improvement in these parameters will result in better survival during the first 24 hours after the injury. Surprisingly, increasing the volume of resuscitation did not improve survival rate in FFP treated rats to an extent similar to WB or HTS treatments. In the latter, mortality was decreased from 38% to 10% and lower.

The dynamic cortical deformation model creates permanent displacement of the cortex and a part of hippocampus. In order to evaluate the extent of the brain damage we performed neurological assessment using the NSS test, which is the animal parallel to Glasgow Coma Scale in humans with an inverse grading system.

The results showed that neurological condition of rats subjected to combined injury was significantly worse in terms of mobility, balance, and hindquarter movement than that of rats subjected to isolated BI. In addition, pathological assessment of cerebral damage showed extended destruction of the perilesional area in CI brains compared to BI brains. Although resuscitation did not result in improved neuronal count and morphology in the perilesional area, improvement in the neurological condition was observed in rats treated with high volumes of blood fluids and low and moderate volumes of HTS, indicating lower extent of secondary injury.

Trauma triggers the immune system, which results in augmented production of cytokines and activation of neutrophils [28]. These activated neutrophils migrate through the circulation, and infiltrate primarily to the lungs, as this is the first capillary bed that receives blood from injured and post-ischemic tissues [29]. In our study, 48 hours after injury, the amount of neutrophils in the blood of rats subjected to isolated or combined injuries was significantly elevated in comparison to sham operated rats. In addition, neutrophil sequestration into the lung was found in all injury groups, and lung tissue of CI rats demonstrated signs of acute lung injury. Resuscitation with moderate or high WB, but not FFP or HTS, reduced the amount of neutrophils in the blood and in the lungs of rats compared to untreated rats.

In our previous study we showed that limiting Ringer’s Lactate resuscitation to 10ml/kg was found beneficial for the rat’s neurological improvement. Here we show that the use of 9ml/kg of hypertonic solution resulted in two-fold reduction of the NSS score. Two clinical trials were performed recently in an attempt to compare HTS and isotonic resuscitation, in patients with severe brain injuries (Glasgow Coma Scale less than 9) [30, 31]. Both studies reported no long-term neurological difference between patients receiving hypertonic versus isotonic in bolus administration. However, the study performed by Bulger et al., included only patients with isolated TBI without hypovolemia [31], which may imply that these patients had milder secondary BI. The study performed by Cooper et al., although enrolled patients with TBI and hypotension, reported the use of colloid resuscitation in addition to HTS resucitation in more than half patients in each group, as a standard protocol of resuscitation [30]. In addition, both studies were performed only in patients with blunt injuries. In our model we could not test the long-term neurological effects on the rats, further than 48 hours after the injury.

Recent studies suggested an early use of FFP in acute coagulopathic patients due to major blood loss (without withholding FFP administration until a certain amount of PRBCs is received) [9, 16, and 17]. However, there is a debate whether early administration of FFP is beneficial to the patients, and whether or not it reduces mortality rates. In 2008, in a retrospective study, Spinella et al., showed that the amount of FFP administration was independently associated with increased survival [32]. A recent meta-analysis, reported that, in patients undergoing massive transfusions, infusion of high FFP: PRBCs ratios was associated with reduction in the risk of death and multiple organ failure (33), however in non-massively transfused patients, FFP infusion was associated with a trend towards increased mortality and increased risk of developing acute lung injury [33, 34]. Our study showed that, although an improvement in hemodynamic, metabolic and behavioral measurements in the group resuscitated with high FFP were similar to the parallel WB resuscitation group, the relatively high mortality rate raises the question whether the use of early FFP may be beneficial.

## Conclusions

Our results indicate that the usage of early FFP, without PRBCs, does not result in improvement of mortality rate or reduction of the brain damage in this model. The HTS however, significantly improved neurological function and mortality rate when used in moderate administration volume (9ml/kg). The best outcome in our study was observed using WB resuscitation. The use of moderate (9ml/kg) or high (18ml/kg) volumes resulted in improved survival, improved neurological outcome and reduced inflammatory response triggered by the combined trauma. The improvement caused by administration of moderate resuscitation volume was comparable to that of the high volume, probably due to similar elevation in hematocrit levels **(S2 Fig)**, thus we conclude that using moderate (9ml/kg) volume for resuscitation is enough to improve the outcome in this model. To summarize, our recommendation for treatment of multiple-trauma patients with brain injury is primarily administration of WB or blood components in ratio of 1:1 when available at pre-hospital setting. In the case when blood components are not available, HTS would be a preferable resuscitation solution in a dose of 9 ml/kg. An important further step in the field of resuscitation fluids in head trauma patients is examination of combinations of PRBCs, FFP and HTS in this model. In addition, our next goal is to test the best resuscitation protocol in a model of penetrating BI combined with uncontrolled HS.

## Supporting information

**S1 Fig. Sodium level in the blood.** Sodium levels were measured from the arterial blood samples at different time-points during the experiment. No changes were observed in control groups **(A)**, and rats resuscitated with whole blood **(C)** and fresh frozen plasma **(D)**. Significant elevation in blood sodium was observed in rats treated with hypertonic saline (HTS) solution: * P<0.05 sham, CI vs. HTS-m, HTS-h. **BI**-Brain Injury, **CHS**-Controlled Hemorrhagic Shock, **CI**-combined injury, l-low (3 ml/kg), m-moderate (9 ml/kg), h-high (18 ml/kg) volume.

**S2 Fig. Blood hematocrit.** Hematocrit was measured from the arterial blood samples at different time-points during the experiment. During hemorrhage, hematocrit was reduced in CHS and CI groups **(A)**, *P<0.05 CI, CHS vs. sham, BI. Administration of hypertonic saline (HTS) **(B)** and fresh frozen plasma (FFP) **(D)** further diluted the blood and lowered hematocrit levels, *P<0.05 CI vs. HTS-m, HTS-h, FFP-m, FFP-h, opposite to whole blood (WB) treatment, which elevated hematocrit levels **(C)** *P<0.05 CI vs. WB-m, WB-h. **BI**-Brain Injury, **CHS**-Controlled Hemorrhagic Shock, CI-combined injury, l-low (3 ml/kg), **m**-moderate (9 ml/kg), **h**-high (18 ml/kg) volume.

